# Transcriptional Regulatory Logic Orchestrating Lymphoid and Myeloid Cell Fate Decisions

**DOI:** 10.1101/2024.11.20.624578

**Authors:** Rasoul Godini, Michaël Chopin

**Author notes:** Corresponding: RG, MC.

## Abstract

The differentiation of hematopoietic stem cells (HSCs) into diverse blood and immune cells is a complex, highly hierarchical process characterized by a series of tightly regulated steps. It involves a sequence of intermediate oligo-potent progenitors making successive binary decisions. This process gradually narrows down lineage possibilities until a final fate is reached. This step-wise process is tightly controlled by Transcription Factors (TFs) and their associated regulome ultimately resulting the differentiation of both lymphoid and myeloid compartments. Here, we set to unravel the lineage-specific gene regulatory circuitry controlling the development of B cells, T cells, Innate Lymphoid Cells (ILCs), and Dendritic Cells (DCs). We employ Weighted Gene Co-expression Network Analysis (WGCNA) to characterize gene modules associated to the lymphoid or myeloid cell fate, enabling the identification of lineage restricted TFs based on their expression patterns. By identifying TFs whose expression is subset-restricted or those with a broader expression in the hematopoietic compartment we construct a regulatory logic that potentially controls the development of these key immune cells. Our results point to conserved regulatory elements between ILCs, Natural Killer cells, and DCs. This analysis unravels an intricate relationship between each cell types and how the expression of key TFs dictate lineage specificity. We particularly dissect the elements associated to conventional DCs and plasmacytoid DCs. In conclusion, our findings shed new lights on regulatory mechanisms controlling blood cell development and offer a blueprint that can be leveraged to better understand the molecular mechanisms underpinning blood cell development.

## Introduction

Haematopoiesis continuously generates billions of blood cells each day, encompassing various cell types with distinct functions essential for maintaining physiological homeostasis and immune competence. In adult mammals, haematopoiesis is driven by hematopoietic stem cells (HSCs) residing in the bone marrow. These HSCs undergo a series of tightly regulated maturation steps, giving rise to successive oligopotent progenitors that progressively loose multipotency to differentiate in a range of specialised mature blood cells [1]. In brief, hematopoietic stem cells (HSCs) are categorized by their self-renewal and multipotency. Long-Term HSCs (LTHSCs) have prolonged self-renewal ability, and give rise to Short-Term HSCs (STHSCs) which possess a more restricted self-renewal potential. STHSCs further differentiate into multipotent progenitors (MPPs), which lack self-renewal ability but contribute to the generation of various blood cell lineages [1]. MPPs subsequently differentiate into common myeloid progenitors (CMPs), producing erythrocytes, megakaryocytes, and monocytes [2], and common lymphoid progenitors (CLPs), which give rise to B cells, T cells, and natural killer (NK) cells [3–6].

This hierarchical process requires the dynamic regulation of TF networks that activate lineage specific gene expression and restrict the differentiation options of haematopoietic progenitors [7–10]. For example, *Runx1* and *Gfi1* are essential for development of HSCs [11, 12]. *Pax5* and *Ebf1* regulate B cells development [13, 14], while PU.1 and IRF8 are instrumental to dictate dendritic cell commitment [15–18]. Thus, the coordinated action of TFs is crucial for directing key regulatory nodes that instruct the emergence of diverse blood cell lineages.

The integration of transcriptome data across various blood cell types, along with chromatin profiling, has significantly advanced our understanding of the regulatory mechanisms governing the haematopoiesis [19, 20]. Adding to that, a comprehensive mapping of the cis-regulatory elements associated with the development of 86 immune cell populations is readily available [21]. Moreover, the role of epigenetic modifications influencing chromatin dynamics during hematopoiesis have been studied [22, 23]. Despite the availability of these resources, there is a critical need to construct detailed gene regulatory networks (GRNs) that elucidate the differentiation, and the functional attributes of the different blood cells produced from HSCs. Such networks are essential for uncovering the overarching principles of gene regulation and the transcriptional circuitry that directs lineage decisions. This understanding will not only offer valuable insights into the process of blood cell formation but also pave the way for exploring novel approaches for ex vivo blood cell production.

Herein we apply network analysis on the high-through transcriptome data of 48 cell types produced by Yoshida and colleague [21] under ImmGen project (www.immgen.org/). Through this analysis, we have identified lineage-specific transcription factors (TFs) and developed a network illustrating their interactions. Additionally, we introduce several novel transcription factors, ranked according to their significance and interaction strength. Lastly, we validate the efficacy of our approach by identifying a novel transcription factor crucial in the decision-making process of a progenitor to differentiate into functionally distinct DC subsets.

## Material and methods

### Data collection

In this study we applied public-accessible datasets GSE109125 of transcriptome data of 127 populations and GSE100738 of ATAC-seq data of 86 primary cell types [21]. The datasets were filtered for stem cells (including LTHSCs, STHSCs, MPP3, MPP4), lymphoid cells (B cells, α/βT cells, and ILCs), and DCs as a group differentiated from both myeloid and lymphoid cells. 91 samples of immune cells were used for further analysis (Supp file 1). For ATAC-seq data we used only the corresponding cells to the transcriptome 39 sample of immune cells (Supp file 2). The ILC group comprises ILC2s, ILC3s and natural killer cells. Additionally, the data was excluded if miss-annotated, and activated T cells and γδ Τ cells were excluded from our analysis. Each lineage encompasses various transitional states. Therefore, analyses were performed through different approaches, including i) lineage-wise that cells are compared as lineage, and ii) subgroup-wise, which compares individual subgroups against each other. For the analysis of the ATAC-seq data, we only focus on the cis-regulatory regions associated with the genes of interest using Open Chromatin Regions (OCRs) data.

### RNA analysis

RNA-seq analyses were performed using edgeR package in R programing language [24]. Samples were grouped as cell lineages and compared to stem cells group (including LTHSC, STHSCs, CLPs and CMPs). A gene was included for the analysis if it has a minimum count of 10 in more than 70% of samples and a total count of 100 across all samples. The genes were significant if log2 fold change of ±0.585 for TFs and ±1 for other genes. Only genes with a false discovery rate (FDR) ≤ 0.05 were considered significant. The specificity of genes to each cell group was calculated as:

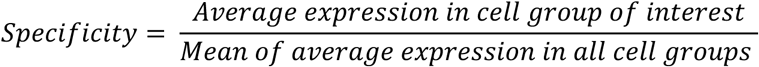

### Gene filtering variable genes and sample clustering

To study the distribution of data, samples were clustered using hierarchical clustering (method = “average”) and Principle Component Analysis (PCA) [25] using R programing language. TFs were identified by comparing the gene list against 1611 from AnimalTFDB 3.0 [26].

### WGCNA analysis

Weighted Gene Co-expression Network Analysis was performed using WGCNA package of R programing language [27]. Based on the clustering, samples were grouped in cell lineages and compared to identify co-expressed genes in each group. For analyzing of all lymphocytes plus DCs, the network was constructed by a soft-power “16” and “signed” type, and modules identified according to a minimum module size of “50” and deep-split of “2”. For analyzing DCs, we used a soft-power “12” due to the smaller number of samples, and deep-split of “2”. The modules were selected based on high association with cell lineages for further analysis. The networks or modules were extracted with a threshold of correlation > 0.01 and thereafter analyzed and visualized using Cytoscape 3.8.2 [28]. Through experiments, genes of interest were filtered using in house developed R scripts.

### Gene regulatory network construction

The regulatory networks were constructed through two approaches. Firstly, by applying WGCNA results through using only the association between TFs and other genes. This approach produces undirected networks of highly expressed genes in particular lineage/s. WGNCA regulatory networks were then combined with information from ChIP Enrichment Analysis from ChEA database [29, 30] and RegNetwork database [31]. Briefly, to obtain potential TF-gene interactions, we submitted a list of cell-specific DEGs (against stem cells) or DEGs present in a particular group of families or all cells, to ENRICHR online tool [29]. We compared the resulted potential regulators and kept ones with a p-value ≤ 0.05 and being specifically expressed in corresponding cell(s). Additionally, the expressing TFs, either DE-TFs or genes detected trough WGCNA, was compared against RegNetwork data [31] to find potential interactions. The final network was constructed by combining all the networks (from WGCNA, ChEA2022, RegNet), and removing duplicate interactions. We constructed a network of only TFs, by presenting only TF-TF interactions and showing the centrality of TFs by adding the number of TF-gene interaction scale as the size of node. The network analysis and adding parameters such as gene expression, gene connectivity within the network, and ATAC-seq data was done using Cytoscape 3.8.2 [28].

### Gene ontology enrichment

DAVID database was used for Gene Ontology (GO) enrichment analysis [32]. The results with a p-values ≤ 0.05 and top matches were used for further analysis.

### BHL15a cloning

Full length coding sequence of flanked by Xho1 sites of *Bhlh15a* was amplified from cDNA generated from RNA isolated from wt splenic pDC. cDNA was cloned into TOPO-TA cloning kit (Invitrogen) according to the manufacturer’s recommendations. *Bhlh15a* cDNA was subsequently subcloned into pMSCV-iresGFP upstream of the IRES-GFP reporter gene.

### Cell culture

Isolated bone marrow cells were cultured in mouse tonicity RPMI-1640 supplemented with 10% heat inactivated fetal calf serum, 2 mM l-Glutamine (Gibco), 50 µM 2-mercaptoethanol (Sigma-Aldrich), and 100 U/ml penicillin/streptomycin (Gibco). 1.5×10^6^ cells/ml were stimulated with 100 ng/ml of Flt3L (Peprotec) for up to 7 days.

### Retroviral infection

Retroviral supernatants were generated by transient transfection of 293T cells with plasmids encoding viral envelope proteins (pMD1-gag-pol and pCAG-Eco), and expression vectors encoding for pMSCV-iresGFP and pMSCV-Bhlh15a-iresGFP using FuGeneHD (Promega). Retroviral supernatants were centrifuged onto RetroNectin (Takara)-coated plates for 45 min at 4000 rpm at 4°C. Cells were then cultivated with the virus in the presence of 4 μg/ml polybrene (Sigma-Aldrich) for 12 hr. 96 h after infection, cells were harvested and stained for flow cytometry analysis.

### Flow cytometry and antibodies

Single-cell suspensions were resuspended in PBS + 2mM EDTA + 0.5% BSA (Sigma-Aldrich) and stained with the indicated antibodies at 4°C. All analyzes were performed on a FACS canto (BD Biosciences) and data were processed using FlowJo. Sorting was performed on a FACSAria (BD Biosciences). Antibodies against CD11c (N418), MHCII (M514.15.2), XCR1 (ZET), SiglecH (551), and CD11b (M1/70) (were purchased from Biolegend). Propidium Iodide (Sigma) was used to exclude dead cells.

### Literature review

To identify known TFs involving in development or functioning of lymphoid cells and DCs, we searched literature with keyword including the name of the TF of interest, and the cell type in which we detected the TFs. We included information in two tables, one for known TFs referencing using PubMed ID (Supp. File 3, S1) and a table showing TFs with no identified function in the corresponding cells. The TFs with no direct study or only having the gene expression in the publication were classified as unknown.

## Results

### High degree of expression similarities across cells of white blood cell lineages

The mammalian blood system consists of multiple lineages stemming for the HSC. To visually capture the development pathways and relationships between the different blood cell types, haematopoiesis is commonly depicted as a tree structure, reflecting the lineage tracing from a common progenitor. In this study, we leveraged publicly available resources to analyse the relationships between progenitors and mature blood cells including B cells, T cells, Innate Lymphocytes and Dendritic Cells [21](Fig 1 a). To identify gene signatures within each group, the cells were clustered into Stem cells, pro-B cells, pre-T cells, B cells, T cells, ILCs, NKs, and DCs. We examined clustering of cells by analyzing transcriptome data using hierarchical and PCA. The results of both methods show that members of each group are clustered together (Fig 1 b and c). Notably, stem cells, proB and preT cells are in very close clusters, suggesting that even as they commit to B or T cells lineages, proB and preT retain a substantial level of stemness (Fig 1 b and c). However, applying a finer resolution by conducting PCA exclusively on the stem cells, proB and preT cells reveals a broader distribution among these cells, thus highlighting sharp differences among these progenitors (Fig 1 b). This analysis revealed that T cells, ILCs, and NK cells are closely grouped demonstrating some degree of similarities between these cells (Fig 1 b). Hierarchical clustering corroborates these findings, aligning closely with the PCA results (Fig 1 c). These analyses show that subsets within each lineage of family exhibit similar transcriptome that set them apart from others. This fact, despite the fine differences within subgroups cells, allows us to compare each subset as a group against stem cells or other lineages for further experiments (Fig 1 a).

**Figure 1.**
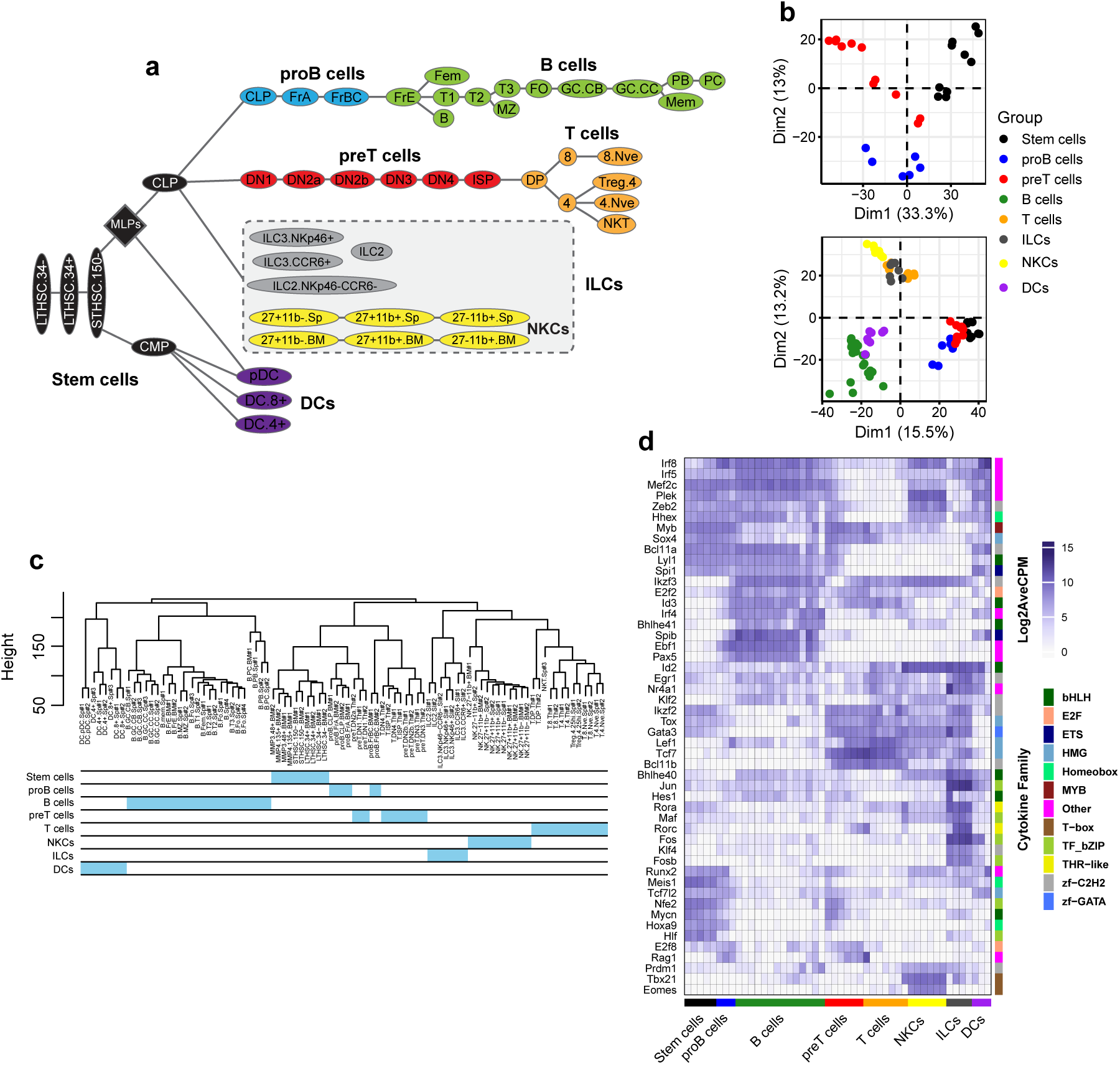
Lymphoid cells and DC lineages classification and gene expression. **A)** The developmental tree of lymphoid cells and DCs, the lineages and subset cell groups used in our study. We consider NKC family as a separate group of ILCs (with modification from [21]). **B)** PCA plots of top 3000 most variable genes of all cell types, and only stem cells to obtain higher resolution. **C)** Hierarchical clustering of all cells. **D)** Heatmap of top 50 most variable TFs across all cell types. Abbreviations: DCs: dendritic cells; ILCs: innate lymphoid cells; NKCs: natural killer cells.

To identify regulatory elements specific to each lineage, we identified the top 50 most variable TFs across all samples. Our results revealed TFs that were expressed either exclusively within a particular lineage or across multiple lineages (Fig 1 d). For example, as expected, *Ebf1* and Pax5 are exclusively expressed in proB and B cells, whereas Spib is also expressed DCs. In contrast, some TFs such as *Ikzf3* and *E2f2* are expressed across many cell types.

The clustering of blood cells into distinct groups reflects high transcriptomic similarity within each group and suggests specific regulatory mechanisms, which we revealed by identifying most variable TFs across cell groups. However, focusing solely on the most variable genes provide an incomplete picture. To enhance our analysis, we performed Weighted Gene Co-expression Network Analysis (WGCNA) approach, which identifies a broader set of group-specific TFs with greater confidence.

### WGCNA Uncovers Cell-Type-Specific Gene Modules and Functions in Immune Cells

WGCNA constructs weighted gene co-expression networks to identify clusters of co-expressed genes and their associations with specific features across samples [27]. We applied this technique to identify genes associated to each cell group. Using soft-power 16 we identified 23 modules correlating to cell groups (Fig 2 a & b). We then focused on lymphoid cells and DCs for further analysis to identify genes associated to each cell type. For each cell type at least one module with the highest correlation was selected. Since B and T cells were strongly correlated to two modules (16 & 5 and 14 & 18, respectively), we included both sets of modules further analysis (Fig 2 b). Each modules comprises a different number of genes, ranging from 121 (Module 26) to 2654 (Module 1). We excluded proB cells and preT because they either resulted in either weakly correlated modules (proB cells) or yielded redundant GO results (preT cells) (data not shown). We then focused on differentiated blood cells that have progressed beyond the stemness stages. To assess the biological relevance of the modules, we performed GO enrichment analysis for genes constructing each module (Fig 2 c). The results indicate that the modules are significantly (p-value ≤ 0.01) correlated to the function or development of the corresponding cell types, as expected. For example, modules 5 and 16 are associated to B cells development (e.g. BCR signaling, antibody production), module 8 is associated with NK cells function (e.g. cytotoxicity, chemokine production) and module 10 is associated with DCs (e.g. inflammatory responses, antigen processing) (Fig 2 c). All modules showed high levels of statistical significance for GO analysis (p-value ≤ 0.01). Notably, module 16 showed very low p-values compared to others, indicating a strong relevance between genes in this module and B cells identity (Fig2 c).

**Figure 2.**
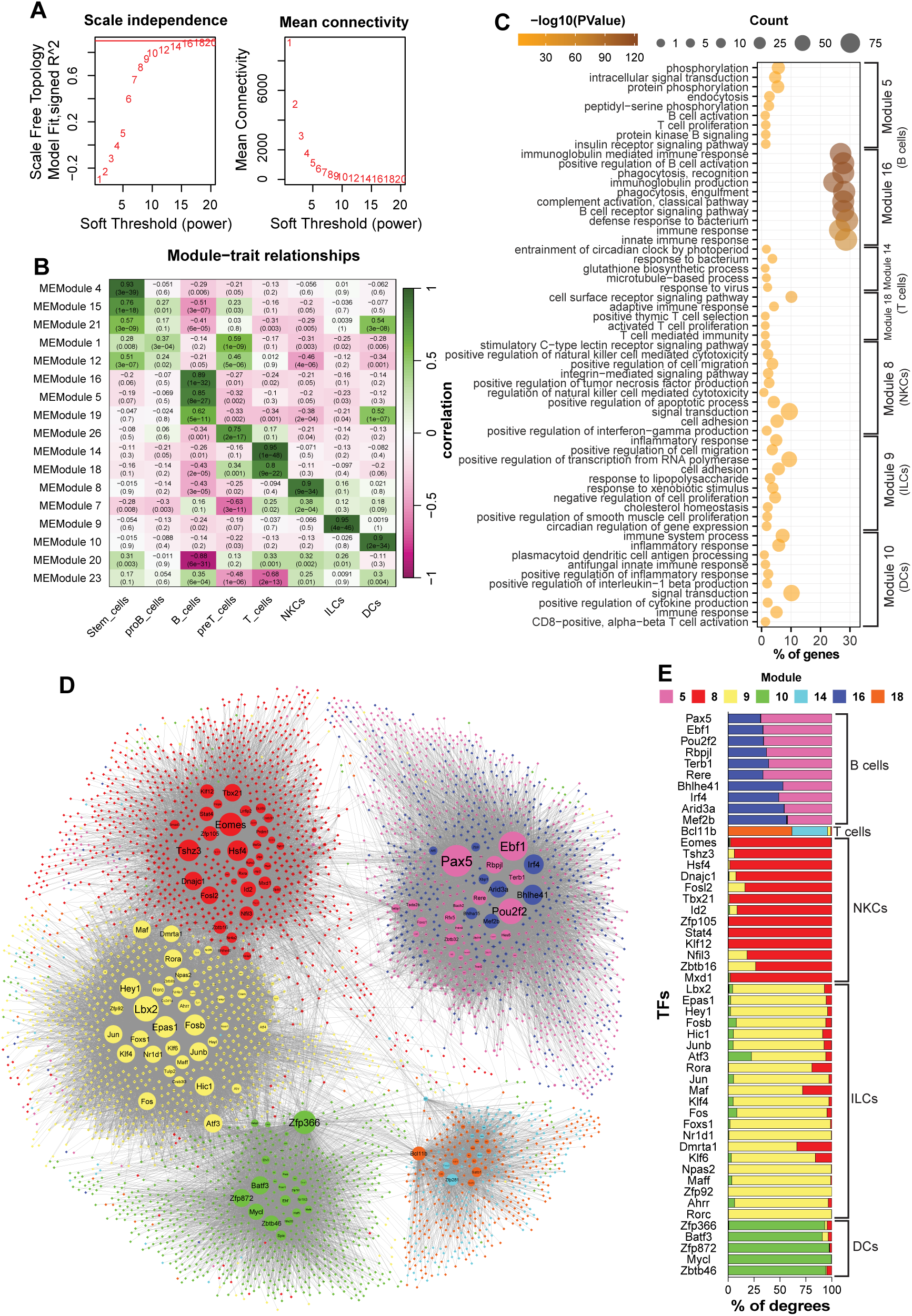
The results of WGCNA and network analysis of the identified modules. **A)** Analyzing of the scale free of the WGCNA. **B)** Heatmap showing the relationships between modules and cell groups. **C)** GO analysis of the related modules. **D)** Integrated network of the related modules to the blood cells. Circles are TFs, and Rhombus are non-TF genes. Larger nodes have higher connectivity (degree). Purple and dark blue: B cells; Cyan and orange: T cells; Red: NKCs; Yellow: ILCs; Green: DCs. **E)** Stacked bar chart shows the distribution of connections with genes to different modules for top 50 most connected TFs. Abbreviations: DCs: dendritic cells; ILCs: innate lymphoid cells; NKCs: natural killer cells.

Altogether, our results showed that WGCNA successfully identified modules correlated to each cell type. We also highlighted that members of each module are associated to cell-specific biological processes, thus indicating the accuracy of our analysis.

### Identifying Cell-Specific Regulatory Elements in Immune Cells

Each module is composed of dozens to hundreds of genes that encode proteins and non-coding RNA molecules involved in different functions. To understand the molecular mechanisms underpinning the formation of each module, we focused our analysis on TFs to identify regulatory elements associated with the development and or function of each cell type. To this end, we retrieved the selected modules (5,8,9,10,14,16 and 18) from the network constructed by WGCNA and applied a filter to select all TFs and their associated genes with a correlation value ≥ 0.02 (weight of edge). The resultant is a network showing intra- and inter-module correlations between TFs and connected genes (Fig 2 d). The topology of the network shows a relatively close interaction between modules of ILCs, NKCs and DCs, in which ILCs and NKCs share several TFs, including *Zbtb16, Nfil3*, *Dmrta1* and *Fosl1* (Fig 2 e). Additionally, Fig 2 d and e show that TFs such as *Atf3*, *Fos*, *Batf3*, and *Zfp366* are correlated to ILCs, NKCs and DCs. Noticeably, in the shared TFs, the majority of connections are intra-modular. In contrast, modules associated to B and T cells show higher isolation. Each one of these cell types is associated to two modules (5 & 16 for B cells; and 14 & 18 for T cells) (Fig 2 b). For each cell type, members of two modules are indistinguishably connected so that TFs are highly connected to the members of both modules. For example, *Bhlhe41*, *Irf4*, and *Mef2b* connections are identically shared between module 5 and 16 (Fig 2 e). Similarly, *Bcl11b* is connected to modules 14 and 18 associated to T cells (Fig 2 e).

In conclusion, our results revealed that ILCs, NKCs, and DCs exhibited partially overlapping co-expression networks and share several TFs. Nonetheless, ILCs and NKCs showed closer association with each other compared to their relationships with DCs which is in line with their respective reported origin in the bone marrow.

### Identifying Cell-Specific Transcription Factors in Immune Cells through Integrated Genomic Analysis

To identify cell-type specific TFs, we examined the identified modules to pinpoint candidates that are either exclusively expressed or expressed at a high level in the corresponding cells. Moreover, the candidate TFs must hold a strong position in their modules, which is defined by module membership (MM) score obtained from WGCNA results. The correlation between MMs and cell types is shown in scatterplots of Fig 3. The scatterplots show TFs and other genes, with corresponding regression lines (*R*) when applicable. We selected TFs with MM ≥ 0.5 to encompass a range of specific (a higher MM score) and more general (a lesser MM score) factors.

**Figure 3.**
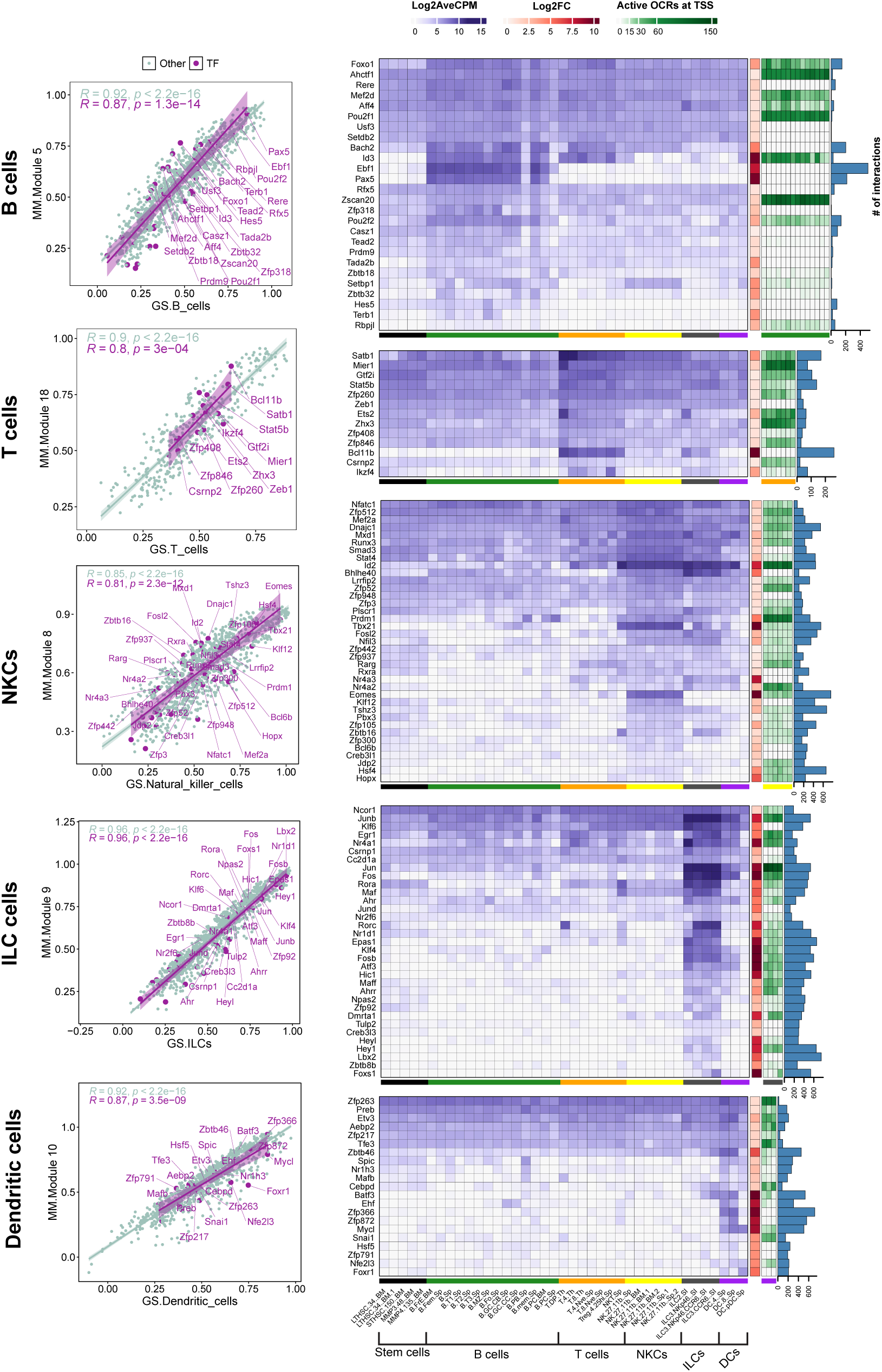
Analysis of TFs in highly related modules to the white blood cells. **Scatter plots:** show the correlation of module membership (MM) and gene significance (GS) of the selected modules. The correlations are shown for TFs and genes belong to other types. Only TFs with a MM ≥ 0.5 are shown. **Heatmaps and bar charts:** The purple heatmaps show the average of gene expression in log2 CPM for selected TFs (MM ≥ 0.05). Orange heatmap shows the changes (Log2FC) in gene expression of corresponding cell families compared to stem cells. A complete list of DEGs in all cell groups is presented in Supp. file 3, S6. Green heatmaps show activity of OCR at the TSS site of the genes in corresponding cells. Blue columns show the degree number of each TF indicating the number of connections with other genes within the module. Black: Stem cells; Green: B cells; Orange: T cells; Yellow: NKCs; dark gray: ILCs; purple: DCs.

In the subsequent step, we characterized candidate TFs by integrating gene expression patterns, chromatin accessibility and their centrality within the modules. We first visualized the expression levels (Log2CPM) and compared these expression levels across all cell types, which provided insights into cell-specify of candidate TFs (Fig 3 purple heatmaps). Additionally, we incorporated expression changes in each cell type relative to stem cells using Log2FC (Fig 3 brown heatmap). Then, we assessed the accessibility of potential cis-regulatory elements at the transcription start site (TSS) of the candidate TFs by incorporating ATAC-seq data specific to corresponding cell types (Fig 3 green heatmap). Finally, we queried the intramodular importance of TFs by analyzing the hubness of the TF in the module, using the degree centrality index, which quantifies the number of connections between a node and all other nodes within the network (Fig 3 bar chart).

Our results showed that number and specificity of TFs expression can vary greatly amongst cell types. Globally, B and T cells showed a limited number of cell-specific TFs compared to other immune cells analysed in our study. For example, Ebf1 and Pax5 are two TFs restricted to B cells, while ILCs exhibit highly expression of several TFs, including *Rorc, Jun, Nr1d1, Fosb, Hic1* and *Epas1* (Fig 3). Interestingly, many of the identified TFs are also expressed across multiple cell types. For example, *Id2* is expressed in NKCs and ILCs, or *Pou2f2* is expressed across all cell types peaking in B cells (Fig 3). These findings suggest that while TFs can function in multiple cell types, their impact may be more pronounced in specific groups, depending on the presence of co-interactors that may play crucial roles in modulating their activity/functionality in each cell type. Concordantly, majority of candidate TFs are also differentially expressed compared to stem cells. Unexpectedly, TFs with cell-specific expression do not always have open chromatin regions (OCRs) at their TSS in the cell type where they are selectively expressed. For example, *Pax5* and *Ebf1* in B cells, *Bcl11b* in T cells, or *Zfp872 i*n DCs have no OCRs at TSS (Fig 3). Finally, the centrality of TFs in modules, is strongly correlated with cell-specify of the expression. For example, *Epas1* and *Lbx2*, which exhibit relatively high and low expression in ILCs, respectively, and almost no expression in the other cells, are each connected to over 600 genes. In contrast, TFs that are expressed in multiple cell types, such as Jun and Fos in ILCs, demonstrate reduced centrality. These findings indicate that the centrality of a TF is associated with the specificity of its expression within a particular cell type.

Altogether, these results have identified multiple cell-specific TFs in white blood cells. We also showed that, in some cases, their expression can be dictated independently of the accessibility of their chromatin at their TSS.

### Regulatory networks controlling lymphocyte development

The distinct patterns of TF expression across blood cells suggest the existence of complex regulatory networks. To better define these networks, we integrated our WGCNA results with data from the RegNetwork [31] and ChEA [30] databases. This approach enabled us to construct a network that integrates multiple aspects of interactions between TFs and other non-TF encoding genes. This includes co-expression, protein-gene and protein-protein interactions. In addition to TFs identified through WGCNA, we included TFs from the ChEA database that are known to regulate genes expressed in each cell type, and any other TFs listed on RegNetwork database. RegNetwork contains gene regulatory networks based on TF, protein-protein interactions, and microRNA information, that are collected and integrated from 25 selected databases [31]. ChEA database is a compilation of chromatin immunoprecipitation (ChIP) studies that identify TF-targeted genes across a range of cell types [30]. In addition to gene clustering derived from the WGCNA results, we also categorized TFs based on their expression level changes compared to the stem cells. Accordingly, TFs could be differentially expressed in a specific (Pax5 in B cells) or multiple cell lineages (Foxo1 and Junb in all cell types). Based on this approach, TFs expressed in multiple cell types were defined as general TFs, either for ILCs, NKCs, and DCs together or all other cells, despite belonging to a specific cell type identified by WGCNA (Fig 4). The resulted network had 3051 nodes 15072 edges (Supp. file 3. S2 and S3, respectively). We analyzed this network and extracted information, such as centrality of the TFs, the type of the connections (WGCNA, ChEA and RegNet) and the cell group(s), to construct a new network restricted to TFs (Fig 4). This network had 170 nodes 467 edges (Supp. file 3. S4 and S5, respectively). Finally, chromatin status was for each connected genes was determined and included as a bar chart showing average OCR activities across all cell types. This visualization provides a comprehensive overview of the regulatory potential of each TF in different cellular contexts.

**Figure 4.**
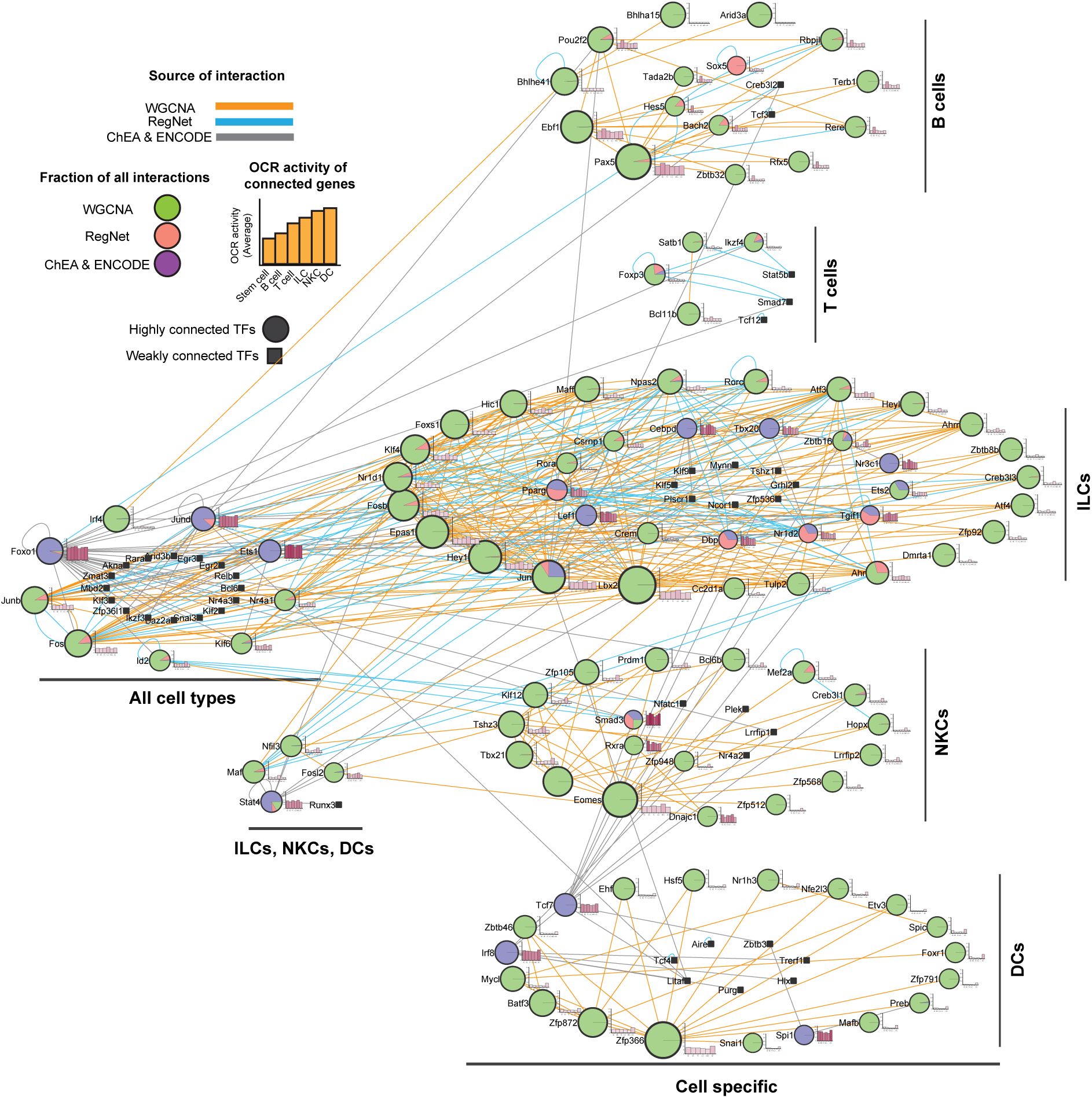
Integrated regulatory network of white blood cells. By combining WGCNA with RegNet and ChEA2022 datasets, we constructed a large network of gene interactions between TF and their non-TF interactors and enriched it with the gene expression data to identify cell specific members expressing in a specific or a group of cells. The resulted network was analyzed for centrality parameters. Next, we construct a smaller network of only TFs Interacting with each other. The TFs centrality calculated in the first network was presented as different in the node size. For each TF the average of OCR activity of its targets across all cell families were calculated and presented as a bar chart next to the nodes. Larger nodes indicate higher degree. Color of the edges show the type on interaction between nodes. The color(s) of the nodes show the fraction of edges identified with data sets. Abbreviations: DCs: dendritic cells; ILCs: innate lymphoid cells; NKCs: natural killer cells.

Using our network, one can predict the significance of TFs in governing blood cell development through factors such as connection type, centrality, specificity, and the activity of its interactors within OCRs. The network shows that WGCNA result is the dominant indicator of the network topology, particularly in individual cell types. In each module, the majority of the TF connections are based on the expression correlation. Hence, having interactions from other sources can enrich WGCNA results. We must notice that TFs within a cluster might have no direct association. For example, there is no detected interaction between Bcl6b and Mef2a in NKCs. In addition, the number of connections show the importance of the TF because larger nodes have higher potential impact on the development. For example, *Pax5*, *Eomes*, and *Foxp3* are known to be important in B cells, NKCs and regulatory T cells development, respectively [33–35]. Specificity is another critical feature for importance of a TF. Based on WGCNA results we identified several cell-specific TFs in each cell type. Remarkably, some of these TFs, *Foxo1*, *Irf4*, *Jund*, *Junb*, and *Fos* are expressed in multiple cell types, suggesting their involvement in controlling the development and or function of these cells, as previously reported by earlier reports. For example, FOXO1 that regulates B cells, T cells and DCs [36–39]. JunB is also involved in development and function of DCs, and NK cells [40–42]. The differential expression of some TFs compared to stem cells, coupled with their shared expression across a range of differentiated immune cells, suggests that these TFs may play critical roles in controlling the development and function of these immune cells, that warrants further investigation.

Finally, we show that the non-TF encoding genes that are interacting (protein-gene interaction or gene co-expression) with the TFs generally have more active OCRs at TSS. For example, genes connected to *Pax5*, *Ebf1*, and *Pou2f2* possess higher active OCRs in B cells compared to other cells (Fig 4). These results indicate a direct correlation between TFs and the accessibility of cis-regulatory elements of their associated genes.

Altogether, our network analysis underscores the importance of several TFs in governing white blood cells development and or function. Additionally, our analysis reveals the interaction and potential roles of TFs within a particular subset of cells or across multiple immune cells. These findings offer valuable insights to study the regulatory mechanisms controlling white blood cells development.

### Identification and Prioritization Strategies for Novel Transcription Factor Candidates Regulating Immune Cell Development and Function

Our analysis identified a range of TFs involved in haematopoiesis, including both well-established TFs and potentially novel candidates. To extract from our analysis, TFs whose function remains unknow in controlling immune cell development, we conducted a comprehensive literature review, focusing on the TFs names and their associated cell families. We classified TFs as known if they have been mechanistically or experimentally validated in relation to blood cell development or function. Using this strategy, we identified 71 novel TFs candidates, whose function in the expressing cells remains to be elucidated (Table 1 and Table 2). Among all TFs 78 (> 50% of all identified TFs) have been already reported to control immune cell development or function (Supp. file 3. S1). Interestingly, all TFs associated with the T cell lineage were all known with reported functions, in contrast ILCs and NKs lineages contain several TFs whose functions remain to be addressed (Supp. file 3. S1). Additionally, our analysis revealed that ubiquitous TFs (found across immune cells) have been mostly experimentally validated at the exception of FOSL2 whose function in ILCs, NKs and DCs remain unknown.

**Table 1.**
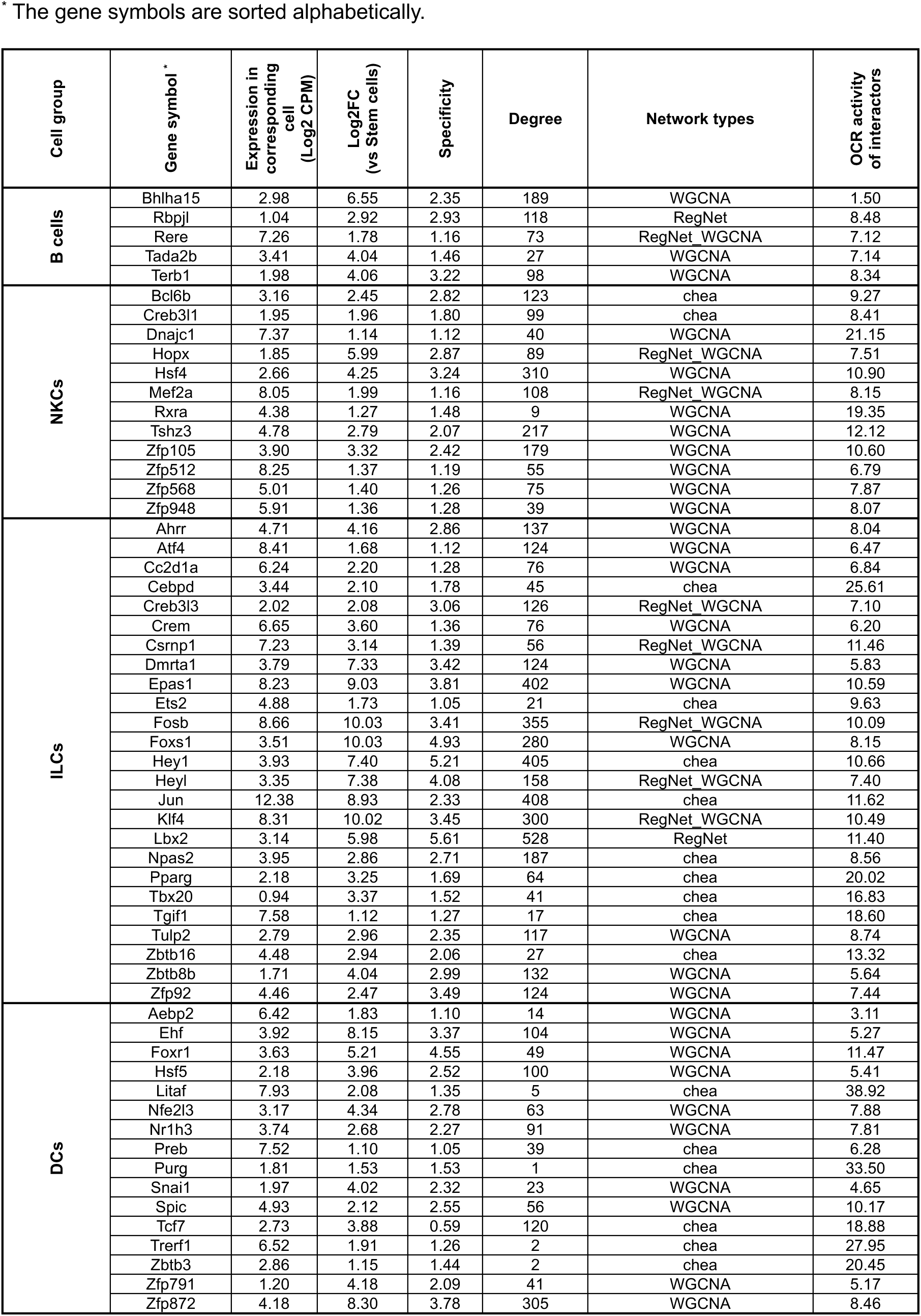
The identified TFs in this study with no reported function.

**Table 2.** The identified TFs associated to DC lineage development with no reported function.

\* The gene symbols are sorted alphabetically.
| Cell group | Gene symbol* | Expression in DC4s (Log2 CPM) | Expression in DC8s (Log2 CPM) | Expression in pDCs (Log2 CPM) | Specificity |
| --- | --- | --- | --- | --- | --- |
| DC4s | E2f2 | 5.88 | 2.82 | 0.09 | 0.83 |
|  | Trps1 | 8.38 | 6.64 | 5.01 | 1.15 |
|  | Zfp263 | 9.43 | 8.39 | 8.01 | 1.08 |
| DC8s | Atf3 | 6.18 | 5.64 | 0.35 | 2.02 |
|  | Ehf | 6.55 | 4.95 | 0.25 | 3.37 |
|  | Fosb | 5.45 | 5.28 | 1.01 | 1.54 |
| pDCs | Bhlha15 | 0.17 | 0.89 | 5.41 | 1.70 |
|  | Cdip1 | 6.00 | 6.01 | 9.00 | 1.04 |
|  | Gcm2 | 0.00 | 0.08 | 3.69 | 2.85 |
|  | Hivep3 | 3.76 | 3.71 | 7.41 | 1.01 |
|  | Sp100 | 7.23 | 6.81 | 8.35 | 0.93 |
|  | Zfp658 | 3.18 | 1.20 | 7.04 | 1.20 |
|  | Zfp810 | 6.06 | 5.97 | 7.86 | 1.10 |

To prioritize the study of unknown TFs, we employed strategies based on expression criteria such as expression levels, fold changes, and cell-specificity. Network centrality metrics were used to rank TFs, with a higher number of connections indicating greater hubness in the regulatory network [43]. Additionally, we integrated data from multiple sources (WGCNA, ChEA, and RegNet) to ensure robust identification. We also applied average OCR activity of TF-associated genes as a sorting criterion, where a higher average suggests a link between TF activity and chromatin accessibility of its target genes (Supp. file 3 S4).

Overall, our analysis validates the effectiveness of our strategy by confirming a substantial number of TFs previously known to regulate the development and function of immune cells. Importantly, our approach also uncovers multiple novel TFs with potential roles in these processes. By employing innovative sorting criteria, we enhance the identification and prioritization of novel TFs involved in white blood cell regulation. To further validate this strategy, we will conduct a case study where in silico analysis led to the identification of a new TF regulating cell fate decisions.

### Deciphering Dendritic Cell Lineage by Unveiling Transcription Factor Network

DCs are antigen presenting cells with the ability to orchestrate adaptive immune responses [44, 45]. Despite their pivotal role in shaping immune surveillance, a comprehensive understanding of their origin and functional specialisation remains a matter of debate [46, 47]. According to the established paradigm of DC specification, myeloid-primed progenitors give rise to both conventional DC subsets (cDCs: type-1 or type-2 cDCs) and plasmacytoid DCs (pDCs). This traditional view serves as the foundation for uncovering the molecular mechanisms that govern DC lineage specification [16, 48]. However, recent studies have challenged this paradigm by disputing the myeloid origin of pDCs, proposing instead that pDCs predominantly originate from a lymphoid-primed progenitor [49–51]. Consistent with this emerging view our findings indicate that, although cDCs and pDCs cluster together, their expression of TFs vastly differs (Fig 3). Therefore, we performed a separate WGCNA analysis for DCs, including stem cells, cDC2s (DC4s), cDC1s (DC8s) and pDCs (Fig 5). Our results showed that stem cells (STHSC, CMP and CLP) are closely clustered, cDC2Ss and CDC1s are grouped together in a separate cluster. In contrast, pDCs form a distinct cluster apart from stem cells and cDCs (Fig 1 a and c), which is reflective of significant differences between conventional and plasmacytoid DCs transcriptomes [21]. We examined the 50 most variable genes in the samples and identified several candidates across cell types. For example, *Relb*, *Id2*, *Fos*, and *Zfp366* are expressed only in cDC2s and cDC1s, not pDCs (Fig 5 b), whereas *Spib* was selectively expressed in pDCs [52]. Conversely, WGCNA results show multiple specific genes for DC types through 4 highly correlated modules (Fig 5 d). We visualized highly correlated TFs (MM > 0.8) in each module to compare their expression levels across cDC2cs, cDC1s and pDCs (Fig 5 e). We found several TFs uniquely correlated with cDC2s (Module 10), including *Zfp263*, *Tsc22d3* and *Rel* (Fig 5 e). In contrast, *Mxd1* and *Zfp872* showed moderate correlation with cDC1s, despite also being slightly expressed by cDC2s. We also searched module 4 that is associated to both cDC2s and cDC1s and identified several highly expressed TFs, such as *Junb*, *Relb*, *Id2*, *Fos*, and *Zfp366* (Fig5 e). Finally, we found multiple TFs highly expressing in pDCs, including the know transcription factors *Tcf4*, *Bcl11a* and *Xbp1*, but also ill-studied TFs such as *Zfp658*, *Cdip1*, *Bhlh15a* and *Hivep3*.

**Figure 5.**
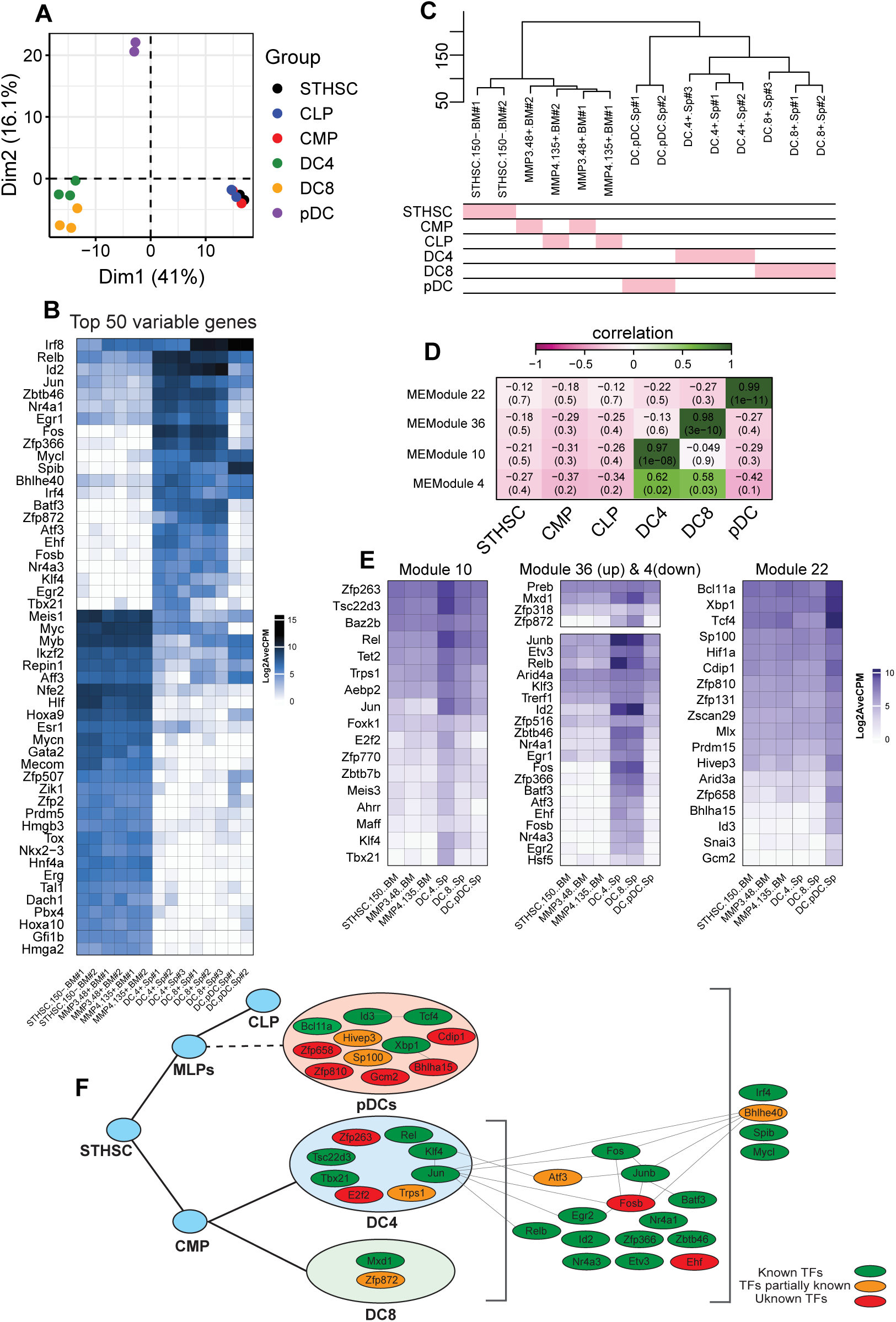
Analyzing of DCs with WGCNA and association of TFs with each cell type. **A)** PCA plot of stem cells and DCs. **B)** Heatmap of top 50 most variable genes in stem cells and DCs. **C)** Hierarchical clustering of stem cells and DCs. **D)** Heatmap of the module and cell relationships of only correlated modules. **E)** Heatmaps of gene expression of TFs with module membership (MM) ≥ 0.5. **F)** The map of TFs expressing in DCs and their identified roles in DCs’ development and/or Function.

Based on these results, we constructed a regulatory logic of TFs that potentially regulate the DC development (Fig 5 f). In this framework, TFs are categorized into pDC, cDC-specific, and general factors that regulate all DCs. Our analysis revealed few known interactions among these TFs. Although a literature review identifies several of these TFs (green nodes) as established key regulators of DC development and function, many candidates (orange and red nodes) remain to be investigated further (Fig 5 d, table 1, and table 2).

### BHLHA15 (Mist1) Inhibits Conventional Dendritic Cell Differentiation

Our data identified several transcription factors (TFs) with poorly defined roles in dendritic cell (DC) differentiation and function (Fig. 5d). Among these, BHLHA15 (also known as Mist1) emerged as a prime candidate for further investigation. BHLHA15 efficiently binds to E-box sequences as a homodimer but can also heterodimers with E-proteins[53]. Importantly the later, particularly E2.2 is crucial in determining the lineage decision between pDCs and cDCs from progenitor cells [48, 54]. This suggests that BHLHA15 may play a significant role in regulating DC lineage fate decisions.

To test this hypothesis, bone marrow progenitors were cultured in the presence of Flt3L and retrovirally transduced with either a control virus encoding GFP or a virus encoding BHLHA15 (Fig 6a). Five days post-transduction, the differentiation of bone marrow-derived DCs was analyzed by flow cytometry. While all three subsets—pDCs (pDCs; SiglecH^+^, CD11b^−^), cDC1s (cDC1s; CD11c^+^ MHCII^+^, XCR1^+^), and cDC2s (cDC2s; CD11c^+^ MHCII^+^, XCR1^−^) were generated from progenitors transduced with either the control or BHLHA15-encoding virus, the overexpression of BHLHA15 significantly impaired the differentiation into conventional DCs, particularly the cDC1 subset (Fig. 6a-b). These findings suggest that BHLHA15 may function as a negative regulator of cDC differentiation.

**Figure 6.**
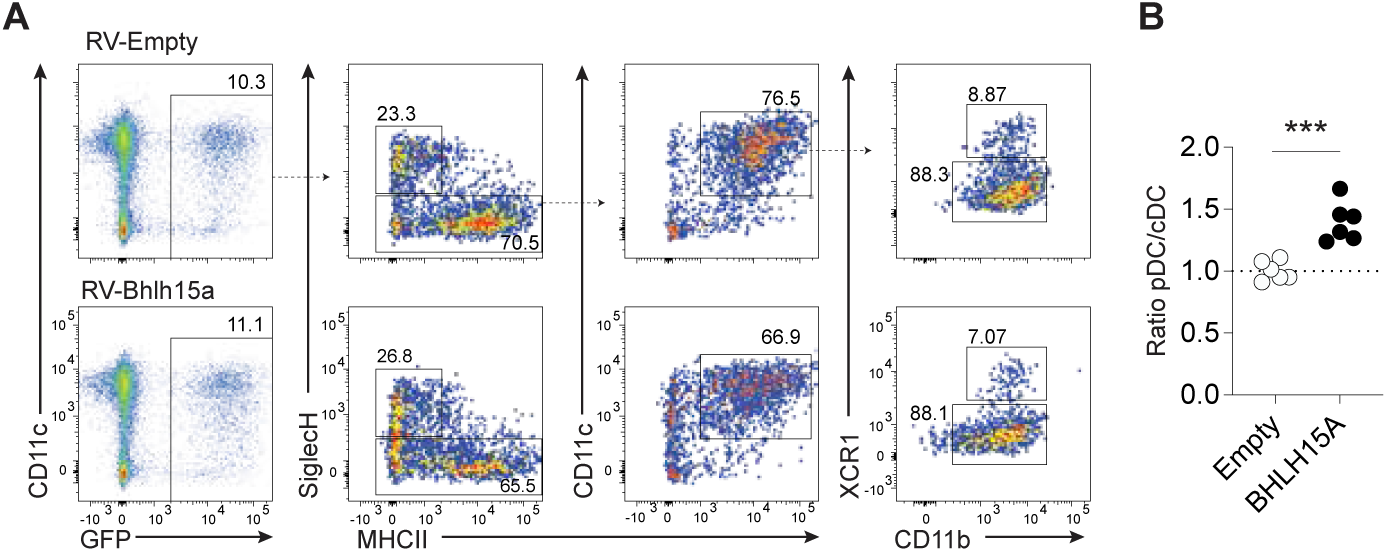
Experimental validation of candidate TFs. **A)** Flow cytometric analysis of bone marrow derived DCs transduced with either GFP encoding (empty: Top row) or BHLHL15a viruses (Bottom row). **B)** The graph represents the ratio of pDC vs cDC (normalised to empty control) of BMDC transduced with control or BHLH15a viruses. The results are shown from two independent experiments (n=3/exp)

## Discussion

A fundamental requirement for a multicellular organism is the establishment of an extensive network of specialized cells, each with unique functions that collaboratively contribute to the development and function of tissues and organs. The hematopoietic system provides a perfect illustration of this concept. For example, multipotent hematopoietic stem cells in the bone marrow undergo differentiation into functionally distinct lineages whose specialization is paramount for vital processes such as oxygen transport, immune surveillance, and haemostasis. As the development of a functional immune system requires dynamic regulation of transcription factor networks that activate lineage specific gene expression and restrict the differentiation options of haematopoietic progenitors, we reasoned that a description of both the wiring and the logic of these transcriptional networks is essential for a complete understanding of immune cell development. Our study leveraged transcriptomic data from highly purified immune cells provided by the ImmGen project, which is well-suited for systems-level analysis. By integrating ATAC-seq data and applying Weighted Gene Co-expression Network Analysis (WGCNA), we constructed a network of co-expressed genes and further enriched it with gene-protein interaction data. This approach enabled us to identify key regulatory circuits and hub TFs associated with the development and function of immune cells.

Network-based analyses have proven effective in uncovering hub genes and proteins across various biological systems [55–60] . Previous studies have investigated regulatory networks in blood cell development, including TF gene regulatory interactions in intestinal ILCs [56] and distinct TF expression patterns in stem and progenitor cells [57]. However, these studies often concentrated on specific cell types or hematopoietic stem cells (HSCs). Although a former study identified the transcriptional regulatory networks for various mouse immune cell types [58], to date most network-based analyses have either focused on HSC or specific immune cell types. In our study we applied WGCNA to identify co-expressed genes uniquely associated with each cell family, providing a detailed network analysis [27]. Our approach benefits from an unbiased analysis that leverages gene expression variability among samples, allowing us to capture insights beyond those provided by just differentially expressed genes (DEGs) [61]. Additionally, we not only identified highly expressed and cell-specific TFs, but also characterized subtle, yet cell-specific, changes in the transcriptome profile. We then enriched the WGCNA results with gene-protein and protein-protein interactions, as well as DEGs, to construct a blueprint regulatory network controlling lymphoid cells and DCs development and function. In addition, applying ATAC-seq demonstrated higher activity of cis-regulatory elements of the TF-associated genes in each group [21]. Altogether, our multi-layered blueprint network highlights central TFs regulating different cell lineages and their functional specificity or redundancy in blood cell development. However, our current network lacks DNA sequence motifs for TF-gene interactions, which would enhance the accuracy of the regulatory network.

Multiple TFs have been identified that regulate blood cell development. The functionality of TFs could be restricted to a specific lineage, such as PAX5 and EBF1 in B cells [13, 14]. In contrast, other TFs, such as FOXO1, have broader roles and influence multiple lineages, including B cells, T cells, and dendritic cells [36–39]. Our study uncovered several TFs with potential roles in lymphoid cells and DCs development and/or function. We combined all related sub-populations as a single lineage due to their close relationships. Notably, proB and preT cells were excluded from analysis of the corresponding matured cells because their characteristics are more closely aligned with those of stem cells. We showed that the expression of most of the identified lineage-specific TFs are highly consistent across all sub-groups and show high statistically significant differential expressions compare to stem cells. These indicate a potential functionality in the cell development and/or function. Noticeably, the lack of chromatin accessibility at the TSS of some of these TFs suggests the existence of regulatory mechanisms through distal-regulatory elements. However, attempts to identify these distal regulatory elements could be compromised by the presence of multiple genes within the100kb region from the TSS, although integration of chromatin accessibility with expression of nearby genes in the relevant cells may help determine if distal regulatory elements are more likely associated with distantly expressed genes rather than nearby non-expressed genes [21].

By performing a separate WGCNA analysis specifically for DCs, our study revealed that cDCs and pDCs possess a set of exclusive regulatory TFs. It further evidenced the central role played by transcription factors such as IRF8, PU.1 (encoded by *Spi1)* and DC-SCRIPT (encoded by *Zfp366)* in controlling cDC identity [17, 18, 62–64]. Interestingly, we identified a substantially higher number of transcription factors associated with cDC2s compared to cDC1s, suggesting not only their functional differences but also that cDC2s may represent a more heterogeneous population of DCs, in contrast to the relatively well-characterized cDC1s in terms of ontogeny and function. [16, 44, 45, 65]. Importantly, our in-silico analysis identified previously uncharacterized transcription factors associated with DC lineage commitment. Among these transcription factors, we demonstrated through a gain-of-function approach that the basic helix-loop-helix transcription factor BHLHA15 plays a regulatory role in progenitor cell fate decisions by inhibiting differentiation into cDCs. While the molecular mechanisms underlying this inhibition are yet to be elucidated, we posit that BHLHA15 expression in progenitors competes with ID2 for E protein binding [53]. This competition could prevent ID2-mediated inhibition of E2.2, thereby blocking the differentiation into conventional dendritic cells (cDCs) [54, 66].

## Conclusion

We developed a comprehensive regulatory network that elucidates the control mechanisms underlying the development and function of immune cells. This network serves as a strategic blueprint for guiding of future research endeavors to decipher the role of newly identified transcription factors in the formation of the immune system.

## Acknowledgement

This work is supported by the National Health and Medical Research Council’s (*NHMRC_1196235*) (MC)

## Author contributions

RG: Conceptualizing, methodology, computational analysis, preparing plots, writing and editing the draft.

MC: Conceptualizing, methodology, laboratory experiments, writing and editing the draft, supervising the project.

## Data availability

Data and scripts will be available upon request.

**Table 1 and Table 2**. Potential TFs in regulating white blood cells development and/or function. The tables present identified TFs in this study with a potential function in regulating development and/or function in corresponding cells. The table of TFs with known functions are present in Supp. file 3 S1.

## Notes

### Competing Interest Statement

The authors have declared no competing interest.

